# Loss of presenilin-1 in smooth muscle cells ameliorates elastin aortopathy

**DOI:** 10.1101/2023.04.15.536809

**Authors:** Junichi Saito, Jui M. Dave, Freddy Duarte Lau, Daniel M. Greif

## Abstract

Smooth muscle cell (SMC) accumulation is central to the pathogenesis of elastin-defective arterial diseases, such as atherosclerosis, pulmonary hypertension and supravalvular aortic stenosis (SVAS). We previously demonstrated that elastin insufficiency activates the Notch pathway in aortic SMCs, resulting in hypermuscularization. Activation of Notch is catalyzed by the enzyme gamma-secretase, but the role of specific catalytic subunits PSEN-1 or PSEN-2 in elastin aortopathy is not defined. This study utilizes genetic approaches to query the role of PSEN-1/2 in the pathogenesis of elastin mutant mice, which model human SVAS. Although endothelial cell-specific *Psen1* deletion does not improve elastin aortopathy, deletion of either *Psen1* in SMCs or *Psen2* globally attenuates Notch downstream gene expression and SMC proliferation, mitigating aortic disease. With SMC-specific *Psen1* deletion in elastin nulls, these rescue effects are more robust and in fact, survival is increased. On the background of *Psen1* deletion in SMCs, global *Psen2* deletion yields additional benefits in regard to elastin aortopathy. Finally, SMC deletion of *Psen1* also attenuates hypermuscularization in newborns heterozygous for the elastin null gene, which genetically mimics SVAS. Taken together, these findings put forth SMC PSEN-1 as a potential therapeutic target in elastin aortopathy.

## Introduction

Defective elastic lamellae and smooth muscle cell (SMC) accumulation are characteristics of diverse arterial diseases, including atherosclerosis, pulmonary hypertension (PH) and supravalvular aortic stenosis (SVAS) (1–5). Although elastin deficiency has been shown to induce SMC hyperproliferation (2, 6), mechanistic links between defective elastic lamellae and SMC hyperproliferation are not well elucidated. Elastin is the major component of circumferential elastic lamellae that alternate with rings of SMCs to form lamellar units in arteries. Heterozygous loss-of-function mutations in elastin gene *ELN* cause SVAS (7, 8). SVAS occurs as an isolated entity or more commonly in Williams-Beuren Syndrome (WBS), which results from a continuous deletion of 26-28 genes on chromosome 7, including *ELN* (9, 10). Elastin mutant mice have been used to model human SVAS (6). *Eln(-/-)* mice exhibit severe aortic phenotypes with increased medial cellularity and lumen obstruction, resulting in early postnatal death (6). In contrast, *Eln(+/-)* mice have modest aortic phenotypes with thinner elastic lamellae and additional SMC layers (11, 12). The only treatment for SVAS is major surgery which carries substantial risk of morbidity and mortality (13), but no pharmacological options are available largely because underlying pathological processes are poorly defined.

The Notch pathway plays integral roles in vascular development and pathology (14–16). Mammals express four Notch receptors, and NOTCH3 is predominantly expressed in arterial SMCs (14, 17). We recently reported that elastin deficiency upregulates the cleaved form of NOTCH3, resulting in aortic hypermuscularization and stenosis (18). Cleavage of Notch receptors is catalyzed by gamma-secretase, which is composed of a catalytic presenilin subunit (PSEN-1 or -2) and accessory subunits (19). Our previous study also demonstrated that systemic pharmacological inhibition of gamma-secretase attenuates elastin aortopathy (18). This finding suggests that gamma-secretase inhibition may be a viable therapeutic strategy for SVAS. However, the gamma-secretase inhibitor used in the prior study is likely to attenuate Notch signaling globally and may have off-target effects. Thus, it is desirable to elucidate the individual roles of PSEN-1 and -2 in elastin aortopathy and gain insights into vascular cell type-specific effects.

PSEN-1 and -2 have been extensively studied in Alzheimer’s disease because mutations in these genes account for ∼90% of the identified mutations in the familial disease (20). Loss of function of *PSEN1* or *PSEN2* causes incomplete digestion of the amyloid β-peptide, which aggregates into neurotoxic amyloid plaques in the brain (21). Conditional *Psen1* deletion in the postnatal forebrain of mice leads to memory impairment (22), whereas global *Psen2* deletion does not have a severe phenotype (23). Mice with compound deletion of forebrain-specific *Psen1* and global *Psen2* exhibit complete loss of PSEN function in excitatory neurons of the postnatal forebrain and gene dosage-dependent cerebral cortex degeneration (24, 25).

Compared to these brain studies, little is known about the role of PSEN-1 and -2 in the cardiovascular system (26). *Psen1(-/-)* mice die shortly after birth with intracranial hemorrhage and bone abnormalities (27), while *Psen2(-/-)* mice are viable and develop mild pulmonary fibrosis (23). Global *Psen1* deletion causes ventricular septal defect, double outlet right ventricle, and pulmonary artery stenosis at embryonic day (E) 12.5-15.5 (28), whereas global knockout of both *Psen1* and *Psen2* induces an un-looped heart at E8.5-9.0 (29). In contrast, combined *Psen1* deletion in adult mice with inducible *CreER^T2^*, which is expressed under the RNA polymerase II promoter, and global *Psen2* deletion protects against angiotensin II-induced cardiac hypertrophy (30). Thus, PSEN1 and PSEN2 play key roles in embryogenesis of the heart and vasculature and pathogenesis of adult cardiovascular disease. However, vascular cell type-specific *Psen1* deletion in any context and the specific roles of PSEN1 and PSEN2 in elastin aortopathy have not been elucidated.

Our current study reports that PSEN-1 and -2 are upregulated in aortic SMCs with elastin insufficiency in human and mouse. Using mouse models, we demonstrate that global *Psen2* deletion partially attenuates aortic hypermuscularization and stenosis in *Eln(-/-)* mutants and *Psen1* deletion in SMCs more significantly mitigates this aortopathy and improves survival. In contrast, *Psen1* deletion in endothelial cells (ECs) does not improve elastin aortopathy. The combination of global *Psen2* deletion and SMC-specific *Psen1* deletion has additive rescue effects on hypermuscularization and stenosis in the elastin-deficient aorta. SMC-specific *Psen1* deletion also rescues hypermuscularization in *Eln(+/-)* mutants. These findings suggest that SMC PSEN-1 plays a main role in elastin aortopathy, providing pre-clinical insight that may facilitate therapeutic strategies to improve efficacy and mitigate off-target effects of gamma-secretase inhibition in this context.

## Results

### Elastin deficiency upregulates presenilin subunits of gamma-secretase

We previously reported that elastin deficiency promotes the cleavage of NOTCH3, resulting in aortic hypermuscularization and stenosis, and that pharmacological inhibition of gamma-secretase attenuates elastin aortopathy (18). Herein, we initially assessed the effect of reduced elastin levels on gamma-secretase subunits PSEN-1 and -2 in human aortic SMCs. Human aortic SMCs were treated with scrambled (Scr) RNA or *ELN*-specific silencing RNA (siRNA) and subjected to quantitative real-time reverse transcription PCR (qRT-PCR) and Western blotting. *ELN* silencing increased transcript and protein levels of PSEN-1 and -2 (Fig. 1A, B). We further analyzed the levels of PSEN-1 and -2 with reduced elastin gene dosage in mice. Aortas were isolated from wild type (WT) or *Eln(-/-)* pups on postnatal day 0.5 (P0.5), and aortic lysates were analyzed by qRT-PCR and Western blotting. Compared to WT, *Eln(-/-)* aortas had higher levels of transcripts (∼6-fold) and protein (∼1.7-fold) for both PSEN-1 and -2 (Fig. 1C, D). Transverse sections of ascending aortas from WT and *Eln(-/-)* pups at P0.5 were stained for PSEN-1 or -2, confirming their upregulation with elastin deficiency (Fig. 1E). These data demonstrate that insufficient elastin in SMCs results in upregulated PSEN-1 and -2.

**Figure 1.**
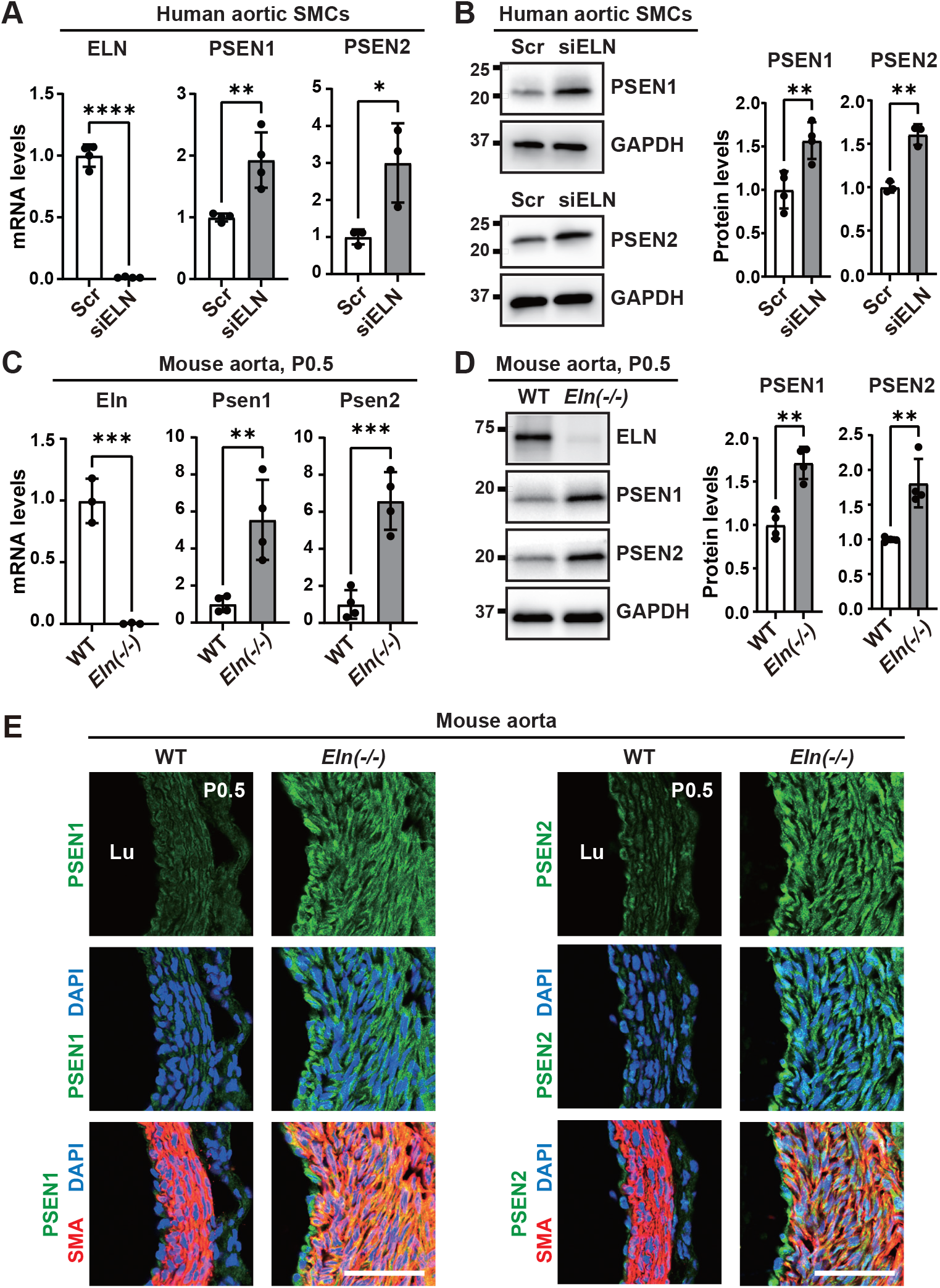
Elastin deficiency upregulates presenilin subunits (PSEN1/2) of gamma-secretase. **(A-D)** Aortic lysates from human aortic SMCs treated with scrambled (Scr) or ELN siRNA (**A, B**) or from wildtype (WT) or *Eln(-/-)* mice at P0.5 (**C, D**) were analyzed. In **A** and **C**, histograms depict transcript levels of ELN, PSEN1, and PSEN2 from qRT-PCR relative to 18S rRNA and normalized to Scr or WT. In **B** and **D**, Western blot and densitometry of protein bands relative to GAPDH is shown. n=3-4. \**P* < 0.05, \*\**P* < 0.01, \*\*\**P* < 0.001, \*\*\*\**P* < 0.0001 vs. Scr or WT by Student’s *t* test. **(E)** Transverse sections of ascending aorta (cranial position, Fig. S1) of WT or *Eln(-/-)* mice at P0.5 stained for α-smooth muscle actin (SMA, SMC marker), nuclei (DAPI), and either PSEN1 or PSEN2. n=5 mice. Lu, lumen. Scale bars, 40 μm.

### Ascending aorta morphology at different cranio-caudal levels in elastin mutants

Similar to SVAS, the hallmark of *Eln(-/-)* newborns is ascending aortic hypermuscularization and stenosis (18, 31); however, a systematic evaluation of the *Eln* null ascending aorta at discrete positions has not been reported. We analyzed serial transverse sections of the ascending aorta from WT or *Eln(-/-)* pups at P0.5 (Fig. S1). We report three positions: 1) cranial – immediately below the aortic arch; 2) middle – between cranial and caudal positions; 3) caudal – where the main pulmonary artery crosses the ascending aorta (Fig. S1A). Our analysis revealed that *Eln(-/-)* ascending aorta at the cranial position has the most significant phenotype with increased medial thickness and area and decreased lumen area compared to WT (Fig. S1B, C). Thus, we analyzed the cranial position of the ascending aorta in all following experiments.

### *Psen2* deletion partially attenuates aortic hypermuscularization and SMC proliferation in elastin mutant aortas

To elucidate the role of PSEN-1 and -2 in elastin aortopathy, we analyzed the effect of *Psen* deletion on the *Eln(-/-)* background. Since *Psen1*(-/-) mice die immediately after birth (27), we started by analyzing *Psen2* global nulls (23). Note the Greif lab received *Psen1(flox/flox)*, *Psen2(-/-)* mice from the Shen lab, and all studies of *Psen2* mutants were performed on the *Psen1(flox/flox)* background. *Eln(+/-), Psen2(+/-)* mice were bred together, pups were harvested at P0.5, and the transverse cryosections of the ascending aorta (cranial position, see Fig. S1) were analyzed. Consistent with our previous study (18), on the *Psen2(+/+)* background, compared to *Eln(+/+)* aorta, *Eln(-/-)* aorta has 2.5-fold increased medial wall thickness and 60% reduction of lumen area (Fig. 2A, B). Global *Psen2* deletion does not alter aortic phenotype on the *Eln(+/+)* background, whereas *Eln(-/-), Psen2(-/-)* mutants had a 23% ± 8% reduction in medial thickness and 158% ± 33% increase in lumen area compared to *Eln(-/-), Psen2(+/+)* mice at P0.5 (Fig. 2A, B).

**Figure 2.**
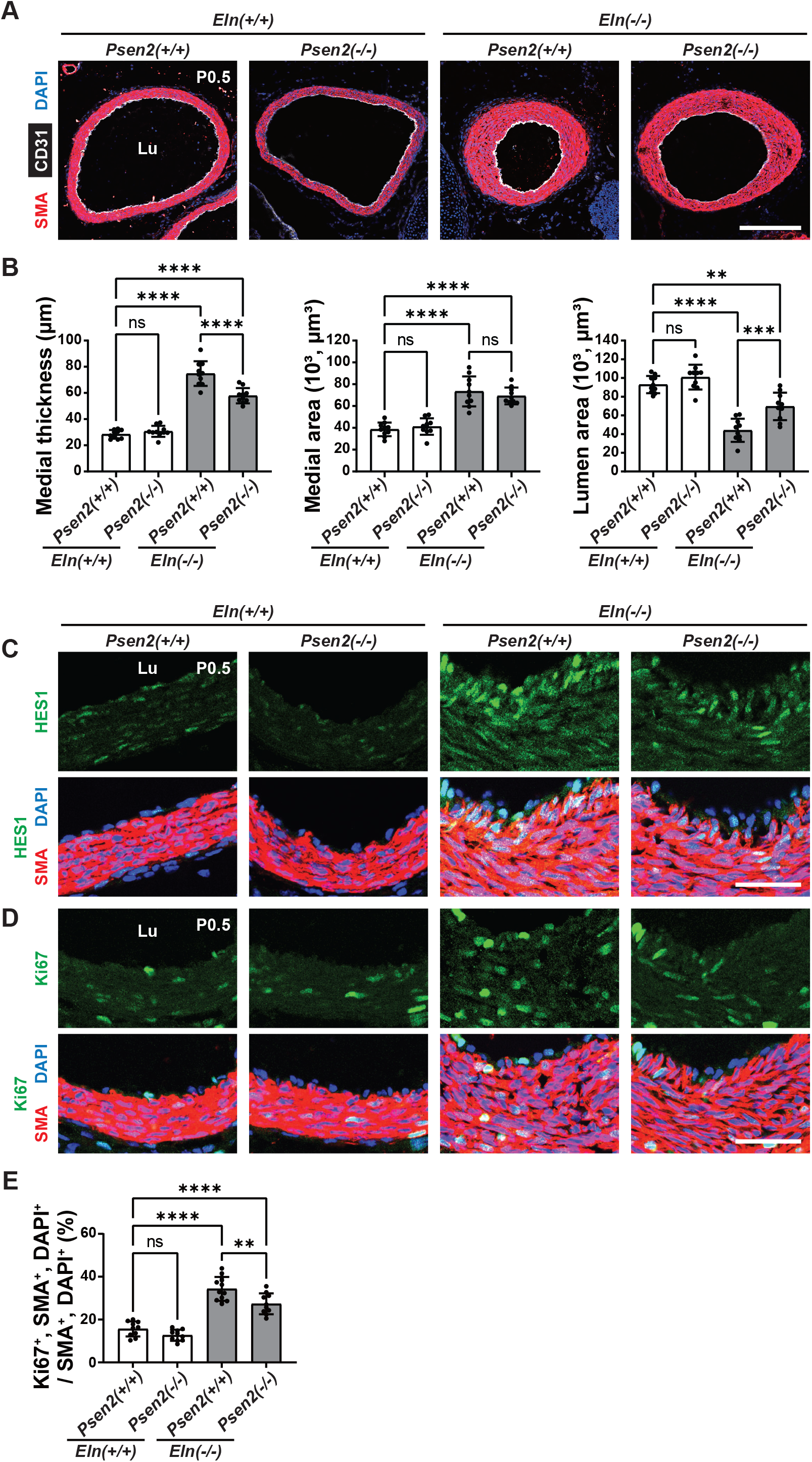
In *Eln* null, *Psen2* deletion attenuates aortic hypermuscularization, stenosis and SMC proliferation. Transverse sections of the ascending aorta (cranial position, Fig. S1) from pups at P0.5 of indicated genotype were analyzed. **(A)** Sections were stained for SMA, CD31 (EC marker), and nuclei (DAPI). **(B)** Histograms represent aortic medial wall thickness, medial area and lumen area from sections as shown in **A**. n=10 mice. **(C, D)** Sections were stained for SMA, nuclei (DAPI), and either HES1 in **C** or Ki67 (marker of proliferation) in **D**. **(E)** Histograms represent percent of SMCs that are Ki67^+^ from sections represented by **D**. n=10-13 mice. Lu, lumen. ns, not significant. \*\**P* < 0.01, \*\*\**P* < 0.001, \*\*\*\**P* < 0.0001 by multifactor ANOVA with Tukey’s *post hoc* test. Scale bars, 200 μm (**A**) and 40 μm (**C, D**).

Immunostains for HES1 (downstream of NOTCH signaling) demonstrate that elastin deficiency upregulates HES1 expression (Fig. 2C). SMCs in the inner part of the medial wall are misaligned in a radial orientation (32), and interestingly, HES1 was most upregulated in this region (Fig. 2C). The expression of HES1 was partially attenuated by *Psen2* deletion (Fig. 2C). Since PSEN-2 is completely abolished in *Psen2(-/-)* aorta (Fig. S2), the remaining expression of HES1 likely reflects PSEN-1 activity. Immunostaining for the proliferation marker Ki67 indicates that *Psen2* deletion modestly reduces elastin deficiency-induced SMC proliferation (Fig. 2D, E). Taken together, *Psen2* global deletion partially attenuates elastin aortopathy.

### *Psen1* deletion with *Acta2-CreER^T2^*, but not *Cdh5-CreER^T2^*, attenuates elastin aortopathy

As global *Psen2* deletion partially rescues elastin aortopathy (Fig. 2) and *Psen1* null mice die immediately after birth (27), we next evaluated the effect of *Psen1* deletion specifically in ECs or SMCs. For investigation of EC PSEN-1, *Psen1(flox/flox)* pups that were also carrying no Cre or inducible *Cdh5-CreER^T2^* and either *Eln(+/+)* or *Eln(-/-)* were analyzed (Fig. S3). We injected pregnant dams at embryonic day (E) 14.5 and E15.5 with 1 mg of tamoxifen and concomitant 0.25 mg of progesterone to minimize the incidence of dystocia (18, 32). At P0.5, newborns were genotyped, and the ascending aortas were sectioned transversely and stained for SMA and CD31. The *Eln(-/-)* aortic phenotype was not altered by EC deletion of *Psen1* (Fig. S4A, B). Immunostaining revealed that *Psen1* deletion in ECs changes neither PSEN1 expression nor Ki67^+^ SMCs in the media (Fig. S4C-F). Thus, EC PSEN1 is not requisite for elastin aortopathy.

To determine the role of PSEN1 in SMCs on elastin aortopathy, *Psen1(flox/flox)* pups that were also carrying no Cre or the inducible *Acta2-CreER^T2^* and either *Eln(+/+)* or *Eln(-/-)* were generated (Fig. 3). Injections of tamoxifen and progesterone at E14.5 and E15.5 (18, 32) led to a robust reduction of PSEN1 in aortic SMCs at P0.5 (Fig. S5). At this time point, the ascending aortas were sectioned transversely and stained for SMA and CD31. In comparison with *Psen1(flox/flox), Eln(+/+)* controls, the medial thickness and area of *Psen1(flox/flox), Eln(-/-)* newborns were increased and the lumen area was decreased. These changes were markedly attenuated in *Acta2-CreER^T2^, Psen1(flox/flox), Eln(-/-)* pups (Fig. 3A, B). Quantitative analysis revealed that on the *Psen1(flox/flox), Eln(-/-)* background, the presence of *Acta2-CreER* leads to a 38% ± 8% reduction in medial thickness, 26% ± 13% reduction in medial area and 225% ± 40% increase in lumen area (Fig. 3B). Notably, in *Eln(+/+)* mice, SMC *Psen1* deletion does not influence aortic morphology. Deletion of *Psen1* in SMCs of elastin nulls leads to decreased medial HES1 expression and Ki67^+^ SMCs (Fig. 3C-F). These findings suggest that SMC *Psen1* is integral to the pathogenesis of elastin aortopathy.

**Figure 3.**
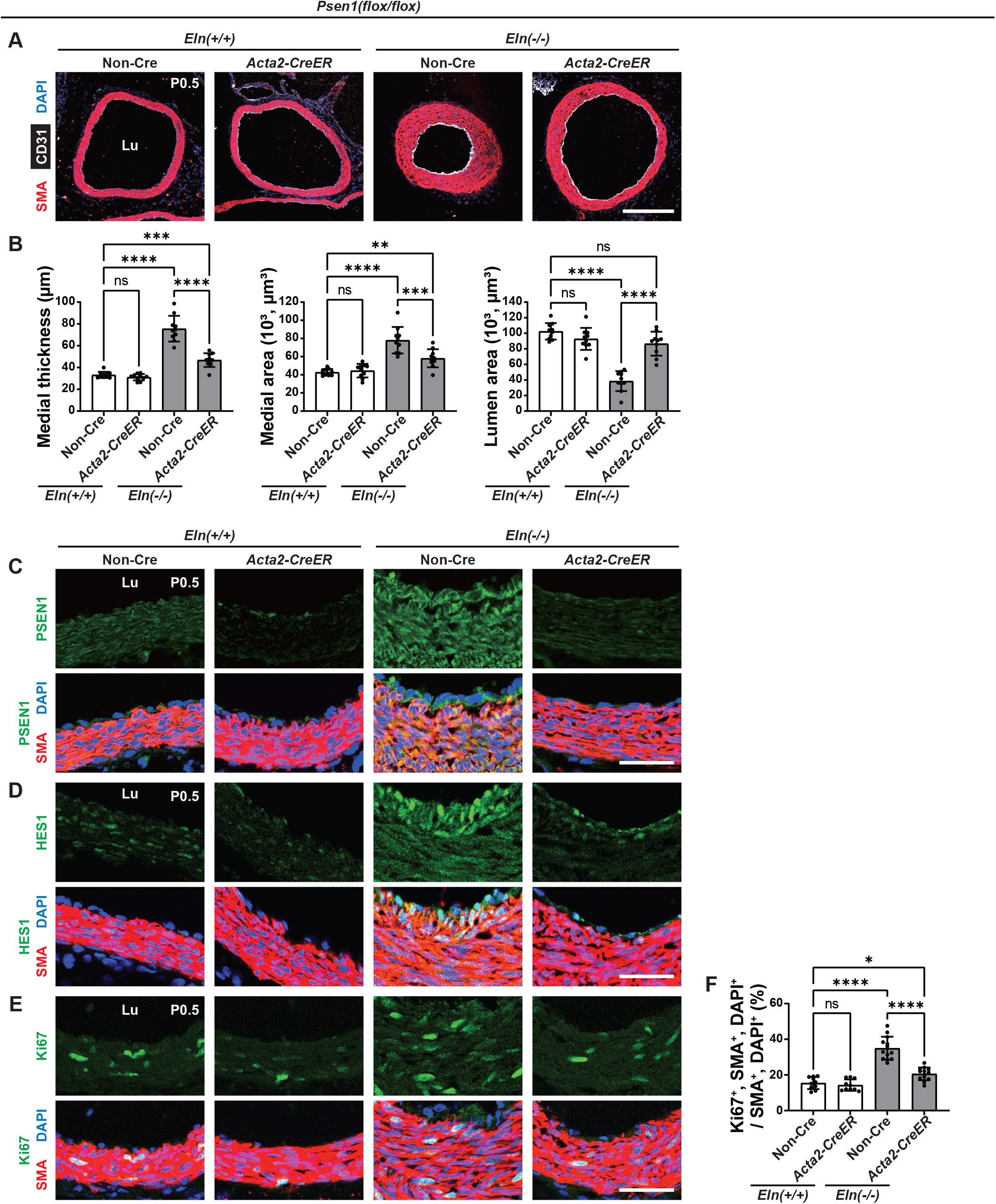
On the *Eln(-/-)* background, SMC-specific *Psen1* deletion reduces hypermuscularization, stenosis and SMC proliferation. Transverse sections of the ascending aorta (cranial position, Fig. S1) from pups at P0.5 of indicated genotype were analyzed. **(A)** Sections were stained for SMA, CD31, and nuclei (DAPI). **(B)** Histograms represent aortic medial wall thickness, medial area and lumen area from sections as in **A**. n=10 mice. **(C-E)** Sections were stained for SMA, nuclei (DAPI), and either PSEN1 in **C**, HES1 in **D**, or proliferation marker Ki67 in **E**. **(F)** Histograms represent percent of SMCs that are Ki67^+^ from sections represented by **E**. n=10-13 mice. Lu, lumen. ns, not significant. \**P* < 0.05, \*\**P* < 0.01, \*\*\**P* < 0.001, \*\*\*\**P* < 0.0001 by multifactor ANOVA with Tukey’s *post hoc* test. Scale bars, 200 μm (**A**) and 40 μm (**C-E**).

### *Psen1* deletion with *Acta2-CreER^T2^* further attenuates aortic hypermuscularization and stenosis in *Psen2(-/-), Eln(-/-)* mice

As a murine pathological model of Alzheimer’s disease, genetic deletion of *Psen1* in the forebrain leads to a compensatory upregulation of *Psen2*, and compound global *Psen2* deletion is required to induce cortical volume reduction (24). Thus, we assessed the combined effect of global *Psen2* deletion and vascular cell type-specific *Psen1* deletion in elastin aortopathy. For EC-specific *Psen1* deletion, *Psen1(flox/flox), Psen2(-/-)* pups that were also carrying no Cre or inducible *Cdh5-CreER^T2^*and either *Eln(+/+)* or *Eln(-/-)* were analyzed (Fig. S6). Pregnant dams were injected at E14.5 and E15.5 with tamoxifen and progesterone as described above (18, 32), and pups were analyzed at P0.5. The aortic phenotype in *Psen2(-/-), Eln(-/-)* pups is not altered by the presence of *Cdh5-CreER^T2^* (Fig. S6A, B). Immunostaining also revealed that expression of PSEN-1, HES1 and Ki67 in the media is not changed by EC-specific *Psen1* deletion (Fig. S6C-F), suggesting that EC PSEN-1 does not have additional rescue effects in *Psen2(-/-), Eln(-/-)* pups.

To determine the role of PSEN-1 in SMCs on *Eln(-/-), Psen2(-/-)* background, a similar experiment was conducted with *Acta2-CreER^T2^*substituting for *Cdh5-CreER^T2^* (Fig. 4). In comparison with *Psen1(flox/flox), Psen2(-/-), Eln(-/-)* newborns, *Acta2-CreER^T2^, Psen1(flox/flox), Psen2(-/-), Eln(-/-)* pups have attenuated medial thickness and area and increased lumen area (Fig. 4A, B). Immunostaining demonstrated marked reduction in expression of PSEN-1 and HES1 and Ki67^+^ SMCs in the media (Fig. 4C-F). These findings indicate additional rescue effects of SMC-specific *Psen1* deletion on *Psen2(-/-)*, *Eln(-/-)* background.

**Figure 4.**
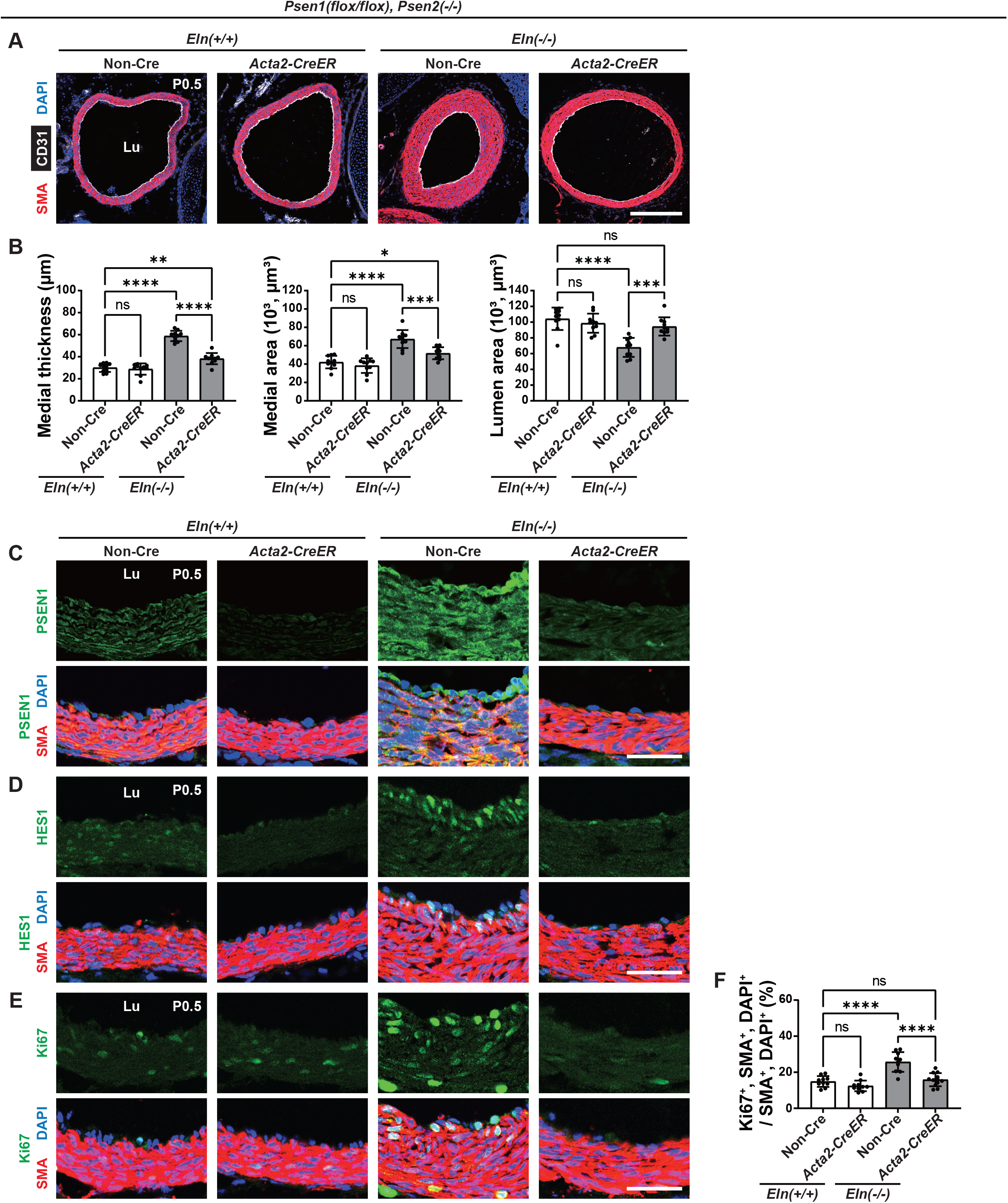
On the *Eln(-/-), Psen2(-/-)* background, SMC-specific *Psen1* deletion further attenuates aortopathy and SMC proliferation. Transverse sections of the ascending aorta (cranial position, Fig. S1) from pups at P0.5 of indicated genotype were analyzed. **(A)** Sections were stained for SMA, CD31 and nuclei (DAPI). **(B)** Histograms represent aortic medial wall thickness, medial area and lumen area from sections as in **A**. n=10 mice. **(C-E)** Transverse sections of the ascending aorta from pups at P0.5 of indicated genotype were stained for SMA, nuclei (DAPI), and either PSEN1 in **C**, HES1 in **D**, or Ki67 in **E**. **(F)** Histograms represent percent of SMCs that are Ki67^+^ in sections represented by **E**. n=10-12 mice. Lu, lumen. ns, not significant. \**P* < 0.05, \*\**P* < 0.01, \*\*\**P* < 0.001, \*\*\*\**P* < 0.0001 by multifactor ANOVA with Tukey’s *post hoc* test. Scale bars, 200 μm (**A**) and 40 μm (**C-E**).

To directly compare effects of global *Psen2* deletion, SMC-specific *Psen1* deletion and the combination, we combined data of Figures 2B, 3B, and 4B in Figure S7. On the elastin null background, SMC-specific *Psen1* deletion is more effective than global *Psen2* deletion in terms of preventing medial thickness and lumen area (Fig. S7). The combination of global *Psen2* and SMC-specific *Psen1* deletion shows the most effective rescue of medial thickness compared with each individual gene deletion (Fig. S7). Quantitative analysis revealed that compound *Acta2-CreER^T2^, Psen1(flox/flox), Psen2(-/-),Eln(-/-)* mutants have a 49% ± 7% reduction in medial thickness, 32% ± 9% reduction in medial wall area and 229% ± 28% increase in lumen area in comparison with *Psen1(flox/flox), Psen2(+/+), Eln(-/-)* mice (Fig. S7).

### Compound *Psen* deletion attenuates aortic tortuosity and extends viability of *Eln(-/-)* mice

Thus far, our findings suggest that combination of SMC *Psen1* and global *Psen2* deletion significantly attenuates the ascending aortic phenotype of *Eln(-/-)* mice. In addition to ascending aorta stenosis, tortuosity of the descending aorta is another hallmark in *Eln(-/-)* mice (33). This tortuosity results from less axial stress and increased SMC proliferation (34). To visualize the descending aorta morphology, yellow latex was injected through the left ventricle at P0.5 (35). Consistent with previous literature (33), *Eln(-/-)* mice have a severely tortuous descending aorta compared to *Eln(-/-)* mice (Fig. 5A) without altering body weight or length (Fig. S8). In elastin WT, neither SMC *Psen1* deletion nor global *Psen2* deletion influences descending aorta tortuosity (Fig. 5A, B). On *Eln(-/-)* background, SMC *Psen1* deletion or global *Psen2* deletion attenuates the tortuosity, and importantly, the combination of these deletions is additive (Fig. 5A, B). Additionally, the survival of elastin null pups is extended by deletion of SMC *Psen1* (Fig. 5C).

**Figure 5.**
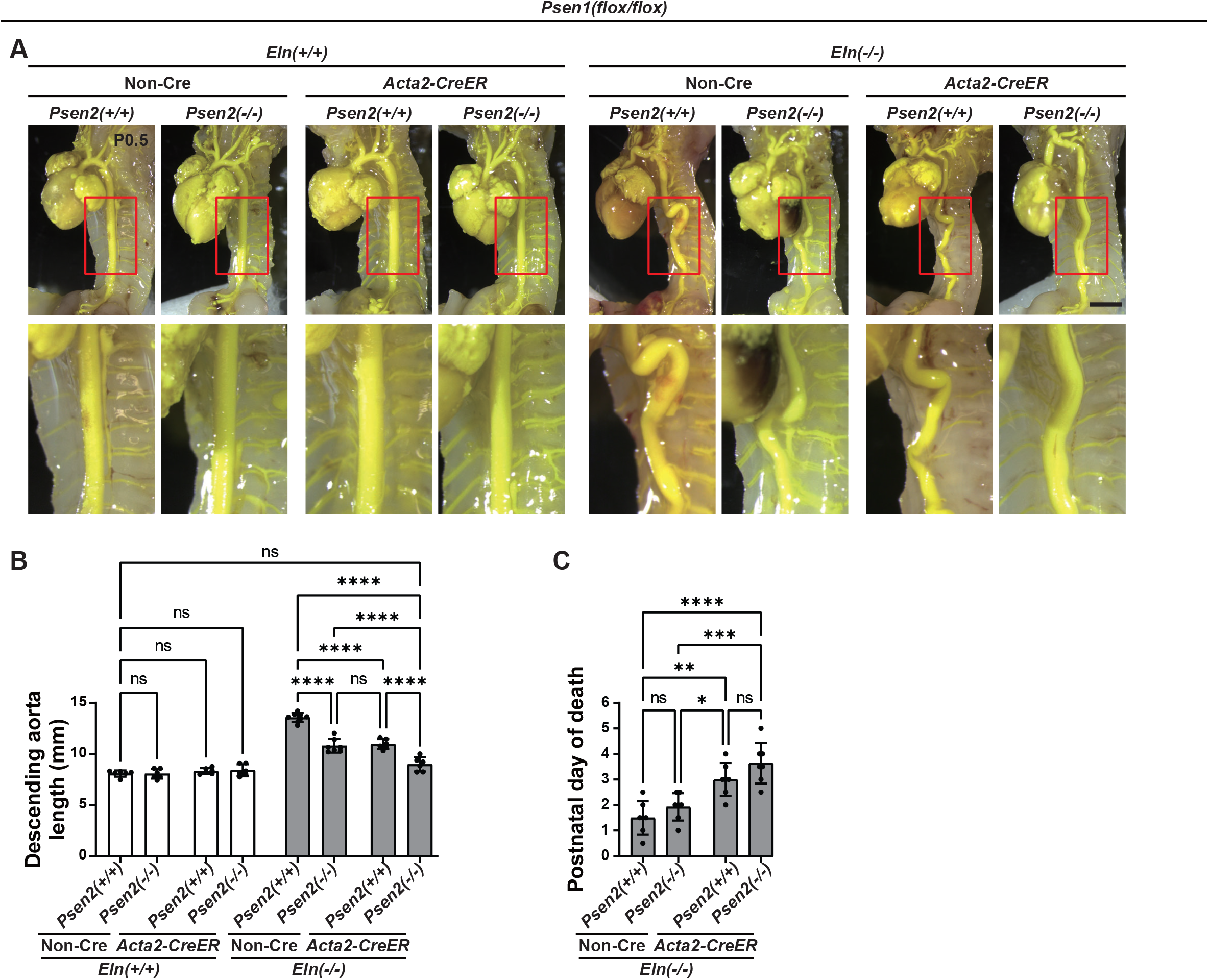
Deletion of *Psen1* in SMCs and *Psen2* globally reduces aortic tortuosity and extends viability of elastin null pups. **(A)** Gross descending aorta morphology was visualized by injecting yellow latex into the left ventricle of newborns of indicated genotypes at P0.5. Lower images are the magnifications of the red boxes. Scale bar, 2 mm. **(B)** Histograms represent the length of the descending aorta from the left brachiocephalic artery to the celiac artery in pups with indicated genotypes at P0.5. n=6-8 mice. **(C)** The age of death of newborns with indicated genotypes is depicted. n=7 mice. ns, not significant. \**P* < 0.05, \*\**P* < 0.01, \*\*\**P* < 0.001, \*\*\*\**P* < 0.0001 by multifactor ANOVA with Tukey’s *post hoc* test.

### Compound of SMC *Psen1* and global *Psen2* deletion attenuates hypermuscularization in *Eln(+/-)* ascending aorta

Finally, we assessed *Eln(+/-)* mice because heterozygous elastin gene mutations cause human SVAS (7). Although *Eln(+/-)* aorta does not have as significant a phenotype as human SVAS, mild hypermuscularization and stenosis are observed in these mice compared to WT mice (Fig. 6). In *Eln(+/-)* mice, SMC *Psen1* deletion, but not global *Psen2* deletion, attenuates hypermuscularization (Fig. 6).

**Figure 6.**
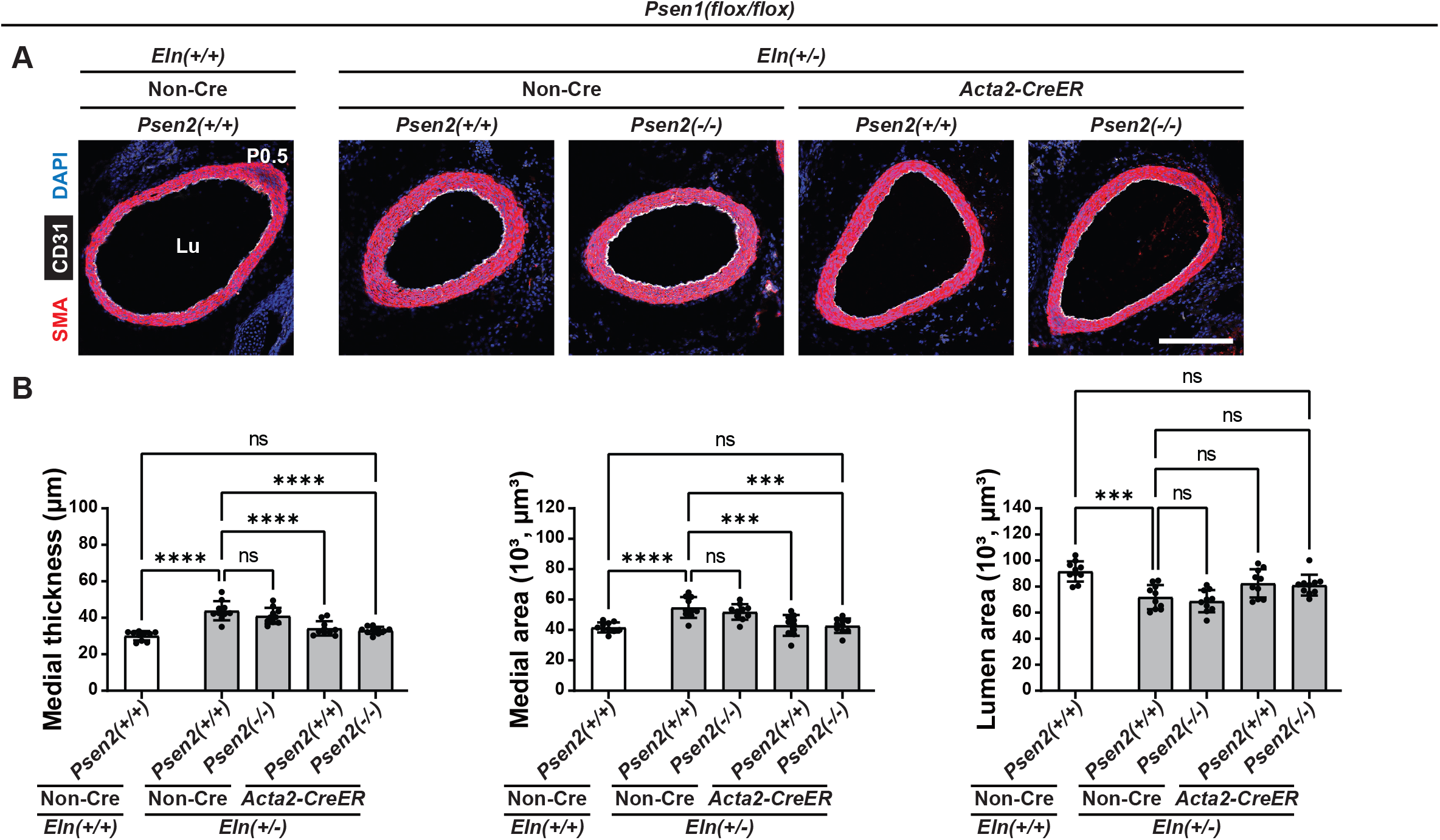
Deletion of *Psen1* in SMCs and *Psen2* globally mitigates hypermuscularization in *Eln(+/-)* mice. **(A)** Transverse sections of the ascending aorta (cranial position, Fig. S1) from pups of indicated genotype at P0.5 were stained for SMA, CD31 and nuclei (DAPI). Lu, lumen. Scale bar, 200 μm. **(B)** Histograms represent aortic medial wall thickness, medial area and lumen area from sections represented by **A**. n=10 mice. ns, not significant. \*\*\**P* < 0.001, \*\*\*\**P* < 0.0001 by multifactor ANOVA with Tukey’s *post hoc* test.

## Discussion

We recently reported that elastin deficiency leads to decreased DNA methylation on promoters of *PSEN-1* and *-2* and increased protein levels of these subunits in human aortic SMCs (18). The current study demonstrates that reduced elastin dosage upregulates mRNA and protein levels of both PSEN-1 and -2 in human and mouse aortic SMCs (Fig. 1) and that the elastin mutant aortic phenotype is rescued substantially by deletion of SMC *Psen1* and partly by global deletion of *Psen2* (Figs. 2-6). Thus, SMC PSEN-1 plays a major role in the pathogenesis of elastin aortopathy.

PSEN-1 and -2 are expressed ubiquitously at comparable levels in most human and mouse tissues (36). They have similar structure, sharing 63% of amino acid residues at identical positions (37, 38); however, PSEN-1 is essential, whereas PSEN-2 is redundant, during development. This difference may derive from distinct subcellular localization patterns of the PSENs. PSEN-1 is present at the plasma and endosomal membranes, whereas PSEN-2 is mainly restricted to the endosomal membrane via an ERTSML motif in the N-terminus (39). Indeed, the higher expression of PSEN1 at the plasma membrane, where Notch receptors are located, may explain why Notch receptor cleavage has been shown to be more dependent on PSEN-1 than PSEN-2 in blastocyst cultures (40). *Psen1(-/-)* pups die at P0.5 with brain hemorrhage and bone abnormalities (27), whereas *Psen2* nulls develop mild pulmonary fibrosis during adulthood (23). The phenotype of *Psen1(-/-)* mice is worsened by *Psen2* deletion as compound *Psen1* and *Psen2* nulls have severe growth retardation at E9.5 and die soon thereafter (23). Our study indicates that ∼70% reduction of PSEN-1 in SMCs markedly improves elastin aortopathy (Figs. 3 and S3), while global *Psen2* nulls have mild effects (Fig. 2).

As catalytic subunits of gamma-secretase, PSEN-1 and -2 activate Notch receptors, which are key players in vascular development and disease (14–16). In addition to proteolytic activity, PSEN-1 is reported to regulate growth and differentiation of endothelial progenitor cells by coupling directly with and inducing degradation of β-catenin (41, 42). However, the roles of PSEN-1 or -2 in definitive ECs or in SMCs have not been elucidated. Our data demonstrate that deletion of *Psen1* and/or *Psen2* is accompanied by reduced Notch downstream signaling (indicated by HES1 expression), suggesting that *Psen1/2* deletion-induced rescue of elastin aortopathy is attributable to suppression of the Notch pathway.

From a clinical perspective, our current and previous studies (18) suggest that gamma-secretase inhibition is a potential therapeutic strategy for SVAS/WBS. Pre-clinical studies of gamma-secretase inhibitors have shown some benefit in cancer (43); however, there are dose-dependent toxicities, including severe diarrhea (44, 45), skin infections (46), and lymphocyte abnormalities (47–49). These adverse effects are due to systemic inhibition of gamma-secretase. As an enzymatic subtype, each gamma-secretase molecule contains one catalytic subunit, PSEN-1 or PSEN-2 (50). The predominant gamma-secretase subtype in human T cell acute lymphoblastic leukemia (T-ALL) contains PSEN-1 (51). In a mouse model of T-ALL, a recently developed PSEN-1 selective inhibitor significantly attenuates disease progression and extends survival and in comparison to a broad-spectrum gamma-secretase inhibitor, reduces adverse effects (51). In the context of elastin aortopathy, SMC PSEN-1 is a major contributor to hypermuscularization, more so than PSEN-2 (Fig. S7). It is also important that SMC-specific *Psen1* deletion reduces hypermuscularization in *Eln(+/-)* mice, whereas global *Psen2* deletion does not (Fig. 6). As human SVAS is caused by heterozygous loss-of-function *ELN* mutations (7), these findings suggest that PSEN-1 selective inhibition may be a promising therapeutic strategy, with reduced adverse effects compared to broad gamma-secretase inhibition, for human SVAS/WBS patients.

Li et al. initially reported that elastin nulls have stenosis of the ascending aorta at the point at which this vessel crosses the pulmonary artery (6), and we previously analyzed the proximal thoracic descending aorta (32) as well as the ascending aorta immediately below the arch (i.e., cranial position as defined herein) (18). To standardize the aortic position for analysis, we analyzed ascending aorta muscularization and lumen size in serial sections in elastin nulls at P0.5. The cranial position of the *Eln(-/-)* ascending aorta shows the most severe phenotype (Fig. S1). Focusing on this position, we found that SMC-specific *Psen1* and/or global *Psen2* deletion significantly attenuates elastin aortopathy (Figs. 2-4). In addition to the stenotic phenotype of the ascending aorta, we analyzed descending aorta tortuosity, another hallmark of elastin nulls (33). To the best of our knowledge, no prior studies have demonstrated rescue of elastin deficiency-induced aortic tortuosity. Our data reveal that deletion of *Psen1/2* ameliorates descending aorta tortuosity (Fig. 5). Since this phenotype is associated with excess SMC proliferation (34), descending aorta tortuosity may be another useful parameter, in addition to ascending aortic stenosis for evaluating effects of genetic/pharmacological interventions in elastin aortopathy.

Reduced gene dosage of SMC *Psen1* alone or of SMC *Psen1* and global *Psen2* in combination extends the survival of *Eln(-/-)* mutants (Fig. 5). Because *Psen* deletion reduces ascending aorta stenosis and descending aorta tortuosity, this increased viability is likely attributable to improved hemodynamics. However, the extension of survival is limited to ∼2 days, probably due to defects in other tissues. Beyond the aortic phenotype, *Eln(-/-)* pups also exhibit severe emphysema (52, 53). SMC-*Psen1* and global *Psen2* deletion are likely insufficient to overcome this strong phenotype. In contrast to *Eln(-/-)* mice, human SVAS/WBS, which are caused by *ELN* haploinsufficiency is not associated with parenchymal lung disease (7, 9, 11). Similar to SVAS/WBS patients, *Eln(+/-)* mice display normal lung development (54), and importantly, deletion of SMC-*Psen1* reduces *Eln(+/-)* aortic phenotypes as well (Fig. 6).

The Notch downstream molecule HES1 is markedly enhanced in the inner media of *Eln(-/-)* aorta, where SMCs exhibit radial orientation (e.g., Fig. 3C) (32). We previously reported that integrin β3 is similarly enhanced in these SMCs (32). Our working model is that the cleaved form of NOTCH3 receptor (NICD3) binds to *Itgb3* promoter and increases expression (18, 32), and thus, the activity of gamma-secretase may be anticipated to be enhanced in the inner media of the *Eln(-/-)* aorta. However, our data demonstrate that PSEN-1 and -2 are upregulated throughout *Eln(-/-)* aortic media, and thus, localized upregulation of HES1 and integrin β3 is not explained by the PSEN-1 and -2 distribution. Although a clear explanation is not yet evident, we speculate on a number of possibilities for HES1 localization. The first possibility for the localized expression of HES1 and integrin β3 in the inner media is the proximity to ECs. EC-secreted factors, including platelet-derived growth factor-B, transforming growth factor-β, and sphingosine-1-phosphate, promote SMC recruitment during embryogenesis (55). Among these factors, shingosine-1-phosphate has been shown to activate the Notch pathway, upregulating HES1 in cancer stem cells (56). In addition, contact between ECs and SMCs leads to SMC differentiation through Notch receptors (57) and connexins (58, 59). Importantly, our previous study demonstrated that the Notch ligand JAG1 is significantly upregulated in ECs of the elastin null aorta, which may activate Notch receptors in SMCs of the inner aortic media (18). Another possible explanation for localized HES1 in the inner layers is the heterogeneous SMC populations in the ascending aortic media. Lineage-tracing experiments of *Eln(+/+)* mice demonstrate that the second heart field (SHF)-derived cells populate the outer media of the ascending aorta, while cardiac neural crest (CNC)-derived cells populate the inner media of the anterior region and contribute transmurally to the media posteriorly (60). A recent study of *Eln* deletion in SHF- or CNC-derived SMCs suggest that SHF-derived SMCs significantly contribute to neointima in the ascending aorta, while CNC-derived SMCs induce medial wall thickening and sporadic neointima (35). This study also performed single cell RNA-seq of *Tagln-Cre, Eln(flox/flox)* ascending aorta and identified SMC subpopulations in the neointima (35). We speculate that select SMC subpopulations with elastin deficiency may potentially promote activity of gamma-secretase, resulting in the localized upregulation of HES1 and integrin β3.

The finding that PSEN-1 in SMCs plays a major role in hypermuscularization in elastin aortopathy may provide insights into the therapeutic strategies for other proliferative vascular diseases, such as pulmonary hypertension (PH). Interestingly, NOTCH3 expression is upregulated in pulmonary arterial SMCs in human and murine PH (61). Furthermore, gamma-secretase inhibition significantly reduces hypoxia-induced PH in mice by attenuating activation of NOTCH3 in SMCs (61). Although the specific catalytic subunit of gamma-secretase involved in PH has not been elucidated, our study suggests that SMC PSEN-1 may play a key role.

In summary, our data reveal that elastin insufficiency in SMCs leads to upregulation of PSEN-1 and -2, which induce Notch signaling and SMC proliferation. Deletion of SMC *Psen1* robustly rescues and global deletion of *Psen2* partly ameliorates aortic hypermuscularization and stenosis of *Eln(-/-)* mice. Thus, inhibiting the gamma-secretase catalytic subunit PSEN-1 in SMCs is an attractive potential therapeutic strategy for SVAS/WBS patients that warrants further investigation.

## Methods

### Animal and tamoxifen treatment

C57BL/6 WT mice were from The Jackson Laboratory. Mouse strains used were *Eln(+/-)* (6), *Acta2-CreER^T2^* (62), *Cdh5-CreER^T2^* (63), *Psen1(flox/flox)* (64), and *Psen2(-/-)* (65). Mice were bred and pups were harvested after birth. As E0.5 was considered the time of vaginal plug, we administered 1 mg tamoxifen (Sigma-Aldrich) with concomitant 0.25 mg progesterone (Sigma-Aldrich) intraperitoneally to pregnant dams on E14.5 and E15.5, and analyzed pups immediately after birth. All mouse experiments were approved by the Institutional Animal Care and Use Committee at Yale University and in accordance with the NIH *Guide for the Care and Use of Laboratory Animals* (National Academies Press, 2011).

### Immunohistochemistry

After euthanasia, pups were fixed in 4% paraformaldehyde for 2 hours. These samples were then washed with PBS for 2 hours and incubated with 30% sucrose in PBS for 24-48 hours at 4°C until the sample sinks. Samples were embedded in OCT compound (Tissue-Tek), frozen on dry ice, and stored at –80°C. Transverse serial cryosections of ascending aortas were cut 8 μm thick starting immediately below the aortic arch and proceeding caudally, ending at the aortic valve. As described in the main text, cranial, caudal, and middle position of ascending aortas were analyzed in Figure S1. All other experiments were performed at the cranial position, which was near the aortic arch. For immunostaining, cryosections were washed with 0.5% Tween 20 in PBS (PBS-T) and incubated with blocking solution (5% goat serum, 0.1% Triton X-100 in PBS) for 1 hour. Then, the sections were incubated with primary antibodies diluted in blocking solution overnight at 4°C. On the next day, sections were washed with PBS-T and incubated with secondary antibodies diluted in blocking solution for 1 hour. Primary antibodies were rabbit anti-PSEN-1 (Cell Signaling Technology, 5643; 1:150), rabbit anti-PSEN-2 (Cell Signaling Technology, 9979; 1:250), rabbit anti-HES1 (Cell Signaling Technology, 11988; 1:1000), rabbit anti-Ki67 (Invitrogen, MA5-14520; 1:250), rat anti-CD31 (BD Pharmingen, 553370; 1:150) and directly conjugated Cy3 mouse anti-SMA (Sigma-Aldrich, C6198; 1:500). Secondary antibodies against rabbit or rat were conjugated to Alexa Fluor 488 or 647 (Molecular Probes, 1:500). DAPI (Sigma-Aldrich, D9542; 1:300) was used for nuclear staining.

### Quantification of aortic morphology

ImageJ software (NIH) was used for quantifications. Transverse sections were used for aortic medial wall and lumen qualifications. As described previously (18), the medial wall thickness from transverse sections of murine aorta were calculated by measuring the distance between the inner aspect of the inner and the outer aspect of the outer SMA^+^ medial layers. Medial and lumen areas were calculated by measuring the area of SMA staining and the area interior to CD31 staining, respectively.

### Cell culture and silencing of *ELN*

Human aortic SMCs (Lonza) were cultured up to passage 6 in M199 medium supplemented with 10% FBS and growth factors (PeproTech, EGF and FGF). For *ELN* silencing, siRNA was transfected as described previously (18). Briefly, human aortic SMCs were transfected with Lipofectamine 2000 (Life Technologies) containing siRNA targeted against *ELN* (Horizon Discovery Biosciences Limited, 50 nM) or Scr RNA for 6 hours. Cells were then washed in M199 medium and cultured for 72 hours prior to collection for qRT-PCR or Western blot analysis.

### qRT-PCR

For pups at P0.5, the entire aorta from the root to the iliac arteries was dissected and was homogenized with tissue homogenizer (Wheaton) in lysis buffer (ThermoFisher). RNA was isolated from mouse aortas and human aortic SMCs with the PureLink RNA Mini Kit (ThermoFisher). Isolated RNA was reverse transcribed with the iScript cDNA Synthesis Kit (Bio-Rad). cDNA was subjected to qRT-PCR on a CFX96 Real-Time System (Bio-Rad) using SsoFast Eva-Green supermix (Bio-Rad) and primers as shown in Supplemental Table 1. mRNA levels were normalized relative to 18S rRNA for cultured cells and to 18S rRNA or Gapdh for murine aortas.

### Western blot

For protein analysis, aortas from pups were mechanically lysed in RIPA buffer with protease and phosphatase inhibitor cocktails (ThermoFisher) on ice with a glass pestle tissue homogenizer (Pyrex). Cultured human aortic SMCs were also collected in the RIPA buffer and vortexed every 10 minutes for 1 hour on ice. Lysates were then centrifuged at 13,000g, 4°C for 5 minutes and supernatants were collected. BCA assay (ThermoFisher) was used to determine the protein concentration. Lysates were prepared in 4× Laemmli sample buffer (Bio-Rad) at 95°C for 5 minutes. Protein samples with Laemmli buffer were resolved by 4–20% SDS-PAGE, transferred to Immobilon PVDF membranes (Millipore), blocked with 5% nonfat dry milk or bovine serum albumin in TBS with tween 20 (TBS-T), washed in TBS-T, and incubated with primary antibodies diluted in blocking solution overnight at 4°C. On the next day, membranes were washed in TBS-T, incubated with HRP-conjugated secondary antibodies (Dako), washed in TBS-T again, and developed with Supersignal West Femto Maximum Sensitivity Substrate (ThermoFisher) on the G:BOX imaging system (Syngene). Western blot analysis used primary antibodies raised in rabbits and targeting PSEN-1 (Cell Signaling Technology, 5643; 1:1000), PSEN-2 (Cell Signaling Technology, 9979; 1:1000), GAPDH (Cell Signaling Technology, 2218; 1:2500), and ELN (raised against exons 6–17 of recombinant mouse tropoelastin; 1:500) (18, 66, 67).

### *Psen1* deletion efficiency

To elucidate *Psen1* deletion efficiency with *Acta2-CreER^T2^*, pregnant dams bearing *Psen1(flox/flox)* embryos with no Cre or *Acta2-CreER^T2^*were injected with tamoxifen at E14.5 and E15.5, and pups were collected at P0.5. As described previously (18), aortas were dissected and the adventitial layer was carefully removed. The remaining medial layer was collected in lysis buffer (ThermoFisher) and mechanically homogenized with a glass pestle tissue homogenizer. RNA was purified for qRT-PCR, and protein was prepared for Western blot as described above. For genotyping, genomic DNA was extracted from the aortic media by incubation with 50 mM NaOH at 95°C for 1 hour, and then 100 mM Tris-HCl pH 7.5 was added to neutralize the digested samples. After centrifugation at 13,000g for 5 minutes, supernatants were used for genotyping PCR with primers as shown in Supplemental Table 2. To determine *Psen1* deletion efficiency with *Cdh5-CreER^T2^*, lung ECs were isolated from *Psen1(flox/flox)* mice with or without *Cdh5-CreER^T2^*at P0.5 using anti-CD31-coated magnetic beads as described previously (18). Isolated ECs were used for qRT-PCR, Western blot, and genotyping.

### Imaging

Fluorescence images of aortic sections were acquired with a confocal microscope (PerkinElmer UltraView Vox Spinning Disc). Adobe Photoshop was used for image processing.

### Vascular casting

After euthanization of pups with isoflurane, the chest wall was opened, and left leg was removed. PBS was injected in the left ventricle and blood was drained through the left iliac artery. Yellow latex (Ward’s Science, 470024-616) was then injected through the left ventricle (35). Mice were then kept moist at 4°C for 3-4 hours to facilitate setting of the injected latex. Mice were subsequently fixed in 4% PFA at 4 °C overnight, followed by fine dissection to demonstrate aortic morphology.

### Statistics

Two-tailed Student’s *t* test and multifactor ANOVA with Tukey’s *post hoc* test were used to analyze data using GraphPad Prism (version 9.4.1). Statistical significance threshold was set at a *P* value of less than 0.05. All data are presented as mean ± SD.

## Author contributions

J.S., J.M.D. and D.M.G. conceived of and designed experiments. J.S. and F.D.L performed them. J.S. and D.M.G. analyzed the results, prepared the figures, and wrote the manuscript. All authors reviewed and provided input on the manuscript.

## Acknowledgments

We thank Dr. Jie Shen at Harvard Medical School for providing *Psen1(flox/flox)* mice and *Psen1(flox/flox), Psen2(-/-)* mice. We also thank Greif laboratory members for their input. Junichi Saito was supported by the JSPS Overseas Research Fellowship from the Japan Society for the Promotion of Science (no. 202260284) and the AHA/CHF Congenital Heart Defect Research Award from the American Heart Association and the Children’s Heart Foundation (23POSTCHF1022933). Funding was also provided by the National Institutes of Health (R35HL150766, R21AG062202, R21NS123469 to D.M.G.), and American Heart Association (Established Investigator Award, 19EIA34660321 to D.M.G.).

---

**Figure S1.**
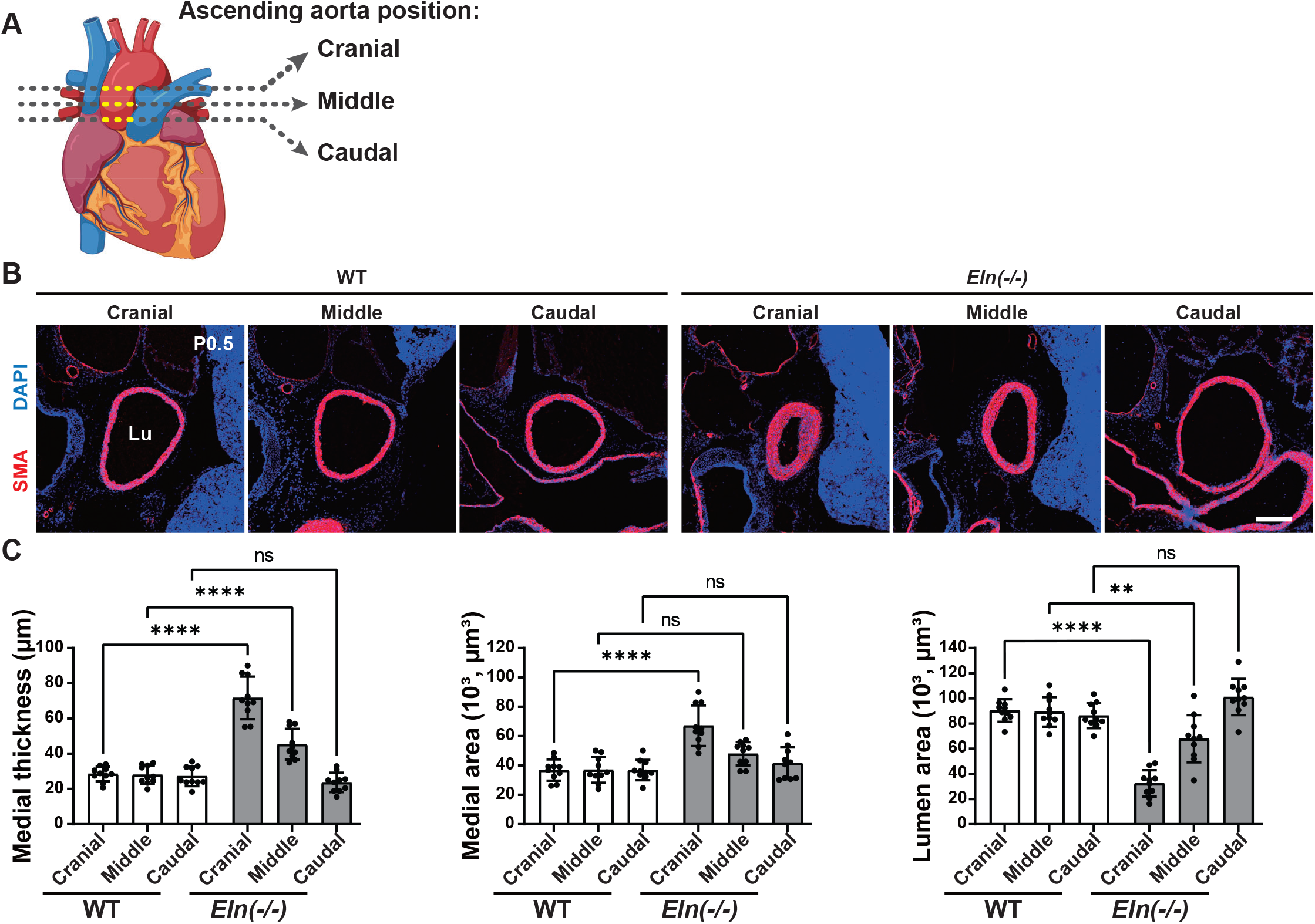
Ascending aorta morphology at different cranio-caudal levels. **(A)** Schematic of the heart and the great vessels. Cranial, middle, and caudal positions of the ascending aorta were analyzed in **B** and **C**. Graphical object was created with BioRender. **(B)** Ascending aortic transverse sections of wildtype (WT) or *Eln(-/-)* mice at P0.5 stained for SMA and nuclei (DAPI). Lu, lumen. Scale bar, 200 μm. **(C)** Histograms represent aortic medial wall thickness, medial area and lumen area from images represented by **B**. n=10 mice. ns, not significant. \*\**P* < 0.01, \*\*\*\**P* < 0.0001 by multifactor ANOVA with Tukey’s *post hoc* test.

**Figure S2.**
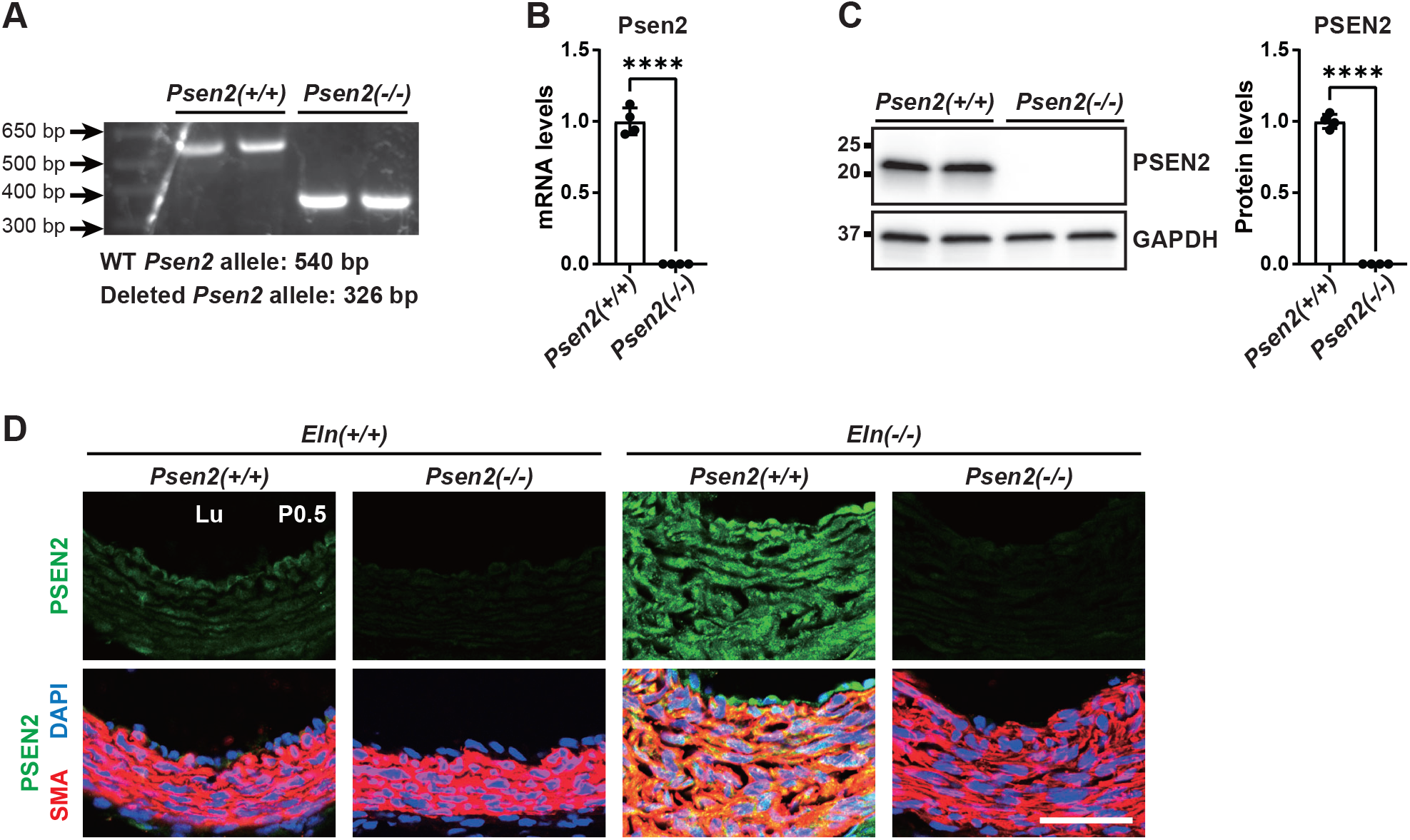
PSEN2 expression in *Psen2* null aorta. **(A)** Extracted genomic DNA from aortas was subjected to PCR with primers targeting *Psen2*. The 540 bp PCR product represents the WT *Psen2* allele, whereas the 326 bp PCR product represents the deleted *Psen2* allele. n=2 mice. **(B)** Isolated RNA from aorta was subjected to qRT-PCR. Histogram represents Psen2 transcript levels relative to 18S. **(C)** Western blot was performed on protein extracted from the aorta. Densitometry of protein bands relative to GAPDH is shown. In **B** and **C**, n=4 mice; \*\*\*\**P* < 0.0001 by Student’s *t* test. **(D)** Transverse sections of the ascending aorta (cranial position, Fig. S1) from pups at P0.5 of indicated genotype were stained for SMA, nuclei (DAPI), and PSEN2. Lu, lumen. Scale bar, 40 μm.

**Figure S3.**
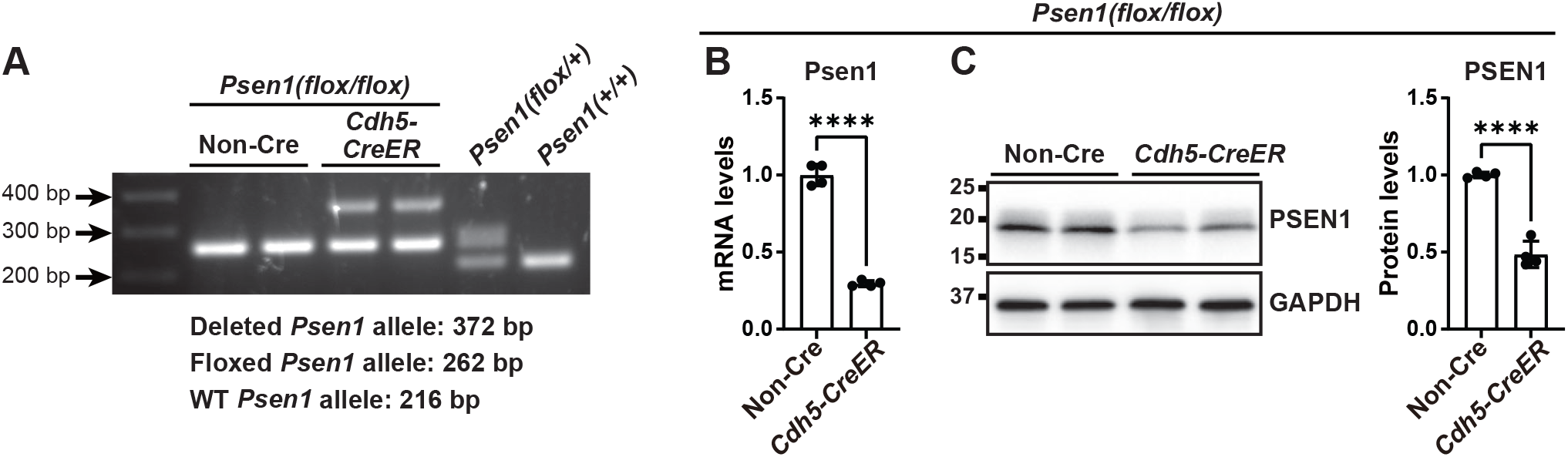
*Psen1* deletion efficiency in ECs of *Cdh5-CreER^T2^*, *Psen1(flox/flox)* mice. Dams pregnant with *Psen1(flox/flox)* embryos also carrying *Cdh5-CreER^T2^*or no Cre were injected with tamoxifen at E14.5 and E15.5. ECs were isolated from lungs of pups at P0.5 and subjected to lysis. **(A)** Extracted genomic DNA was subjected to PCR with primers flanking *Psen1*. The 216 bp PCR product represents the WT *Psen1* allele, whereas the 262 bp fragment represents the floxed *Psen1* allele. The 372 bp PCR product represents the deleted *Psen1* allele (68). **(B)** Isolated RNA was reverse transcribed and subjected to qRT-PCR. Histogram represents Psen1 transcript levels relative to 18S rRNA. **(C)** Western blot was performed on protein extracted from isolated ECs. Densitometry of protein bands relative to GAPDH is shown. n=4 mice. \*\*\*\**P* < 0.0001 by Student’s *t* test.

**Figure S4.**
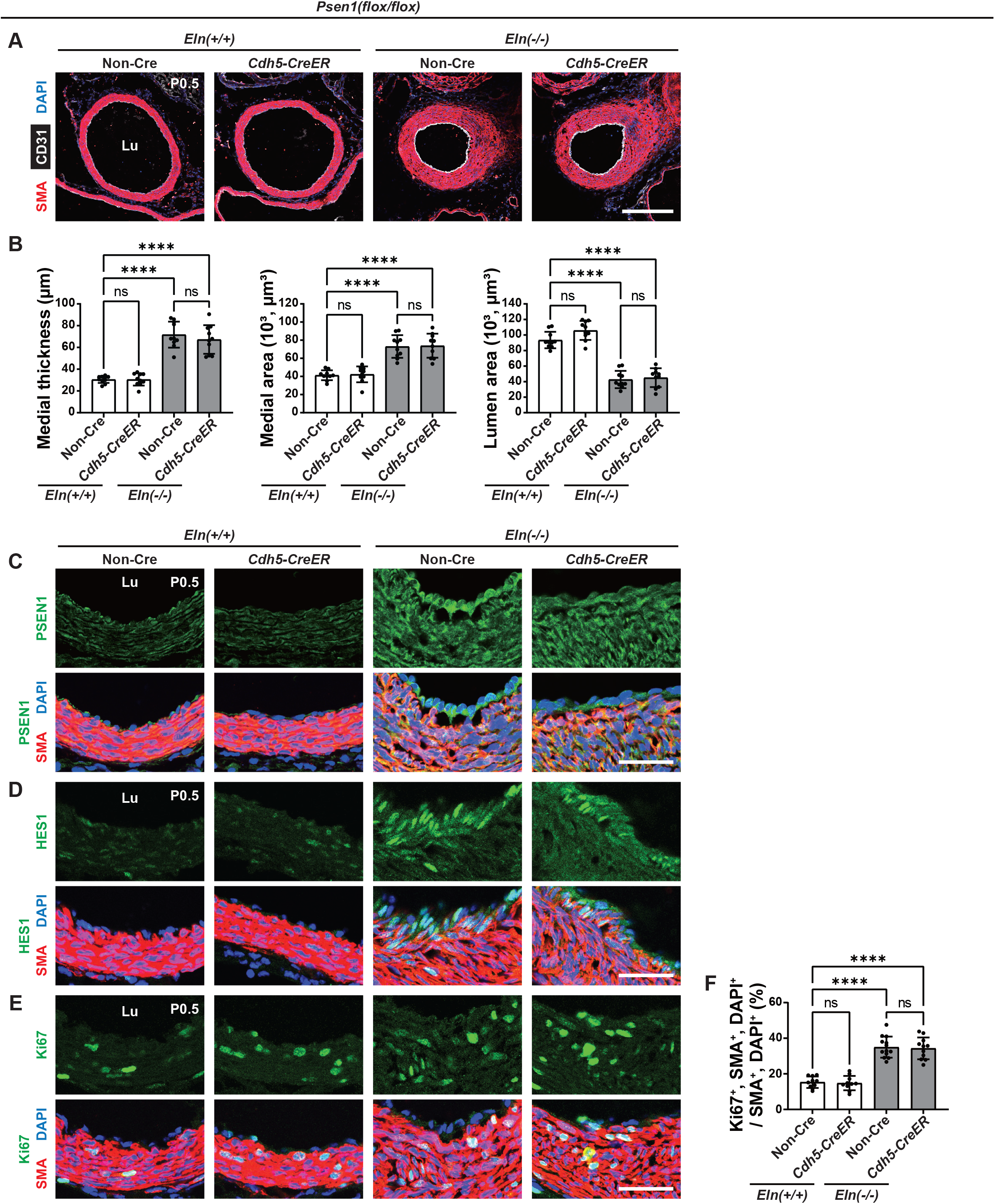
On the *Eln(-/-)* background, EC-specific *Psen1* deletion does not reduce hypermuscularization and stenosis. Transverse sections of the ascending aorta (cranial position, Fig. S1) from pups at P0.5 of indicated genotype were analyzed. **(A)** Sections were stained for SMA, CD31, and nuclei (DAPI). **(B)** Histograms represent aortic medial wall thickness, medial area and lumen area from sections as in **A**. n=10 mice. **(C-E)** Sections were stained for SMA, nuclei (DAPI), and either PSEN1 in **C**, HES1 in **D**, or Ki67 in **E**. **(F)** Histograms represent percent of SMCs that are Ki67^+^ in sections represented by **E**. n=10-13 mice. Lu, lumen. ns, not significant. \*\*\*\**P* < 0.0001 by multifactor ANOVA with Tukey’s *post hoc* test. Scale bars, 200 μm (**A**) and 40 μm (**C-E**).

**Figure S5.**
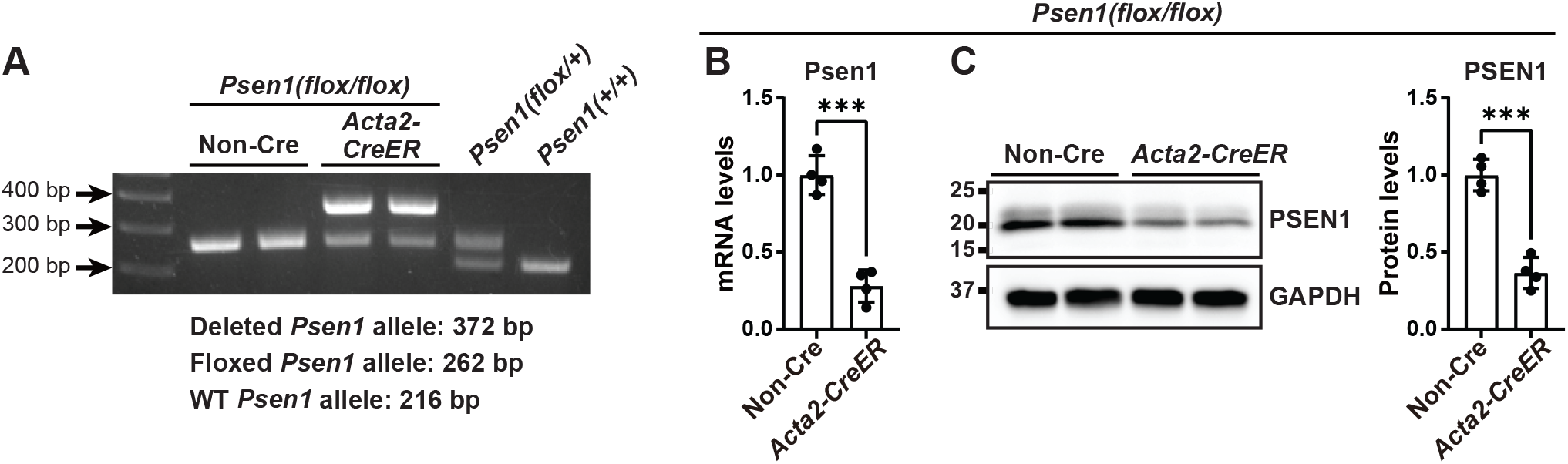
*Psen1* deletion efficiency in aortic SMCs of *Acta2-CreER^T2^*, *Psen1(flox/flox)* mice. Dams pregnant with *Psen1(flox/flox)* embryos also carrying *Acta2-CreER^T2^* or no Cre were injected with tamoxifen at E14.5 and E15.5. Aortas were isolated from pups at P0.5 and subjected to lysis after adventitia removal. **(A)** Extracted genomic DNA was subjected to PCR with primers flanking *Psen1*. The 216 bp PCR product represents the WT *Psen1* allele, whereas the 262 bp fragment represents the floxed *Psen1* allele. The 372 bp PCR product represents the deleted *Psen1* allele (68). **(B)** Isolated RNA was reverse transcribed and subjected to qRT-PCR. Histogram represents Psen1 transcript levels relative to 18S rRNA. **(C)** Western blot was performed on protein extracted from the aorta. Densitometry of protein bands relative to GAPDH is shown. n=4 mice. \*\*\**P* < 0.001 by Student’s *t* test.

**Figure S6.**
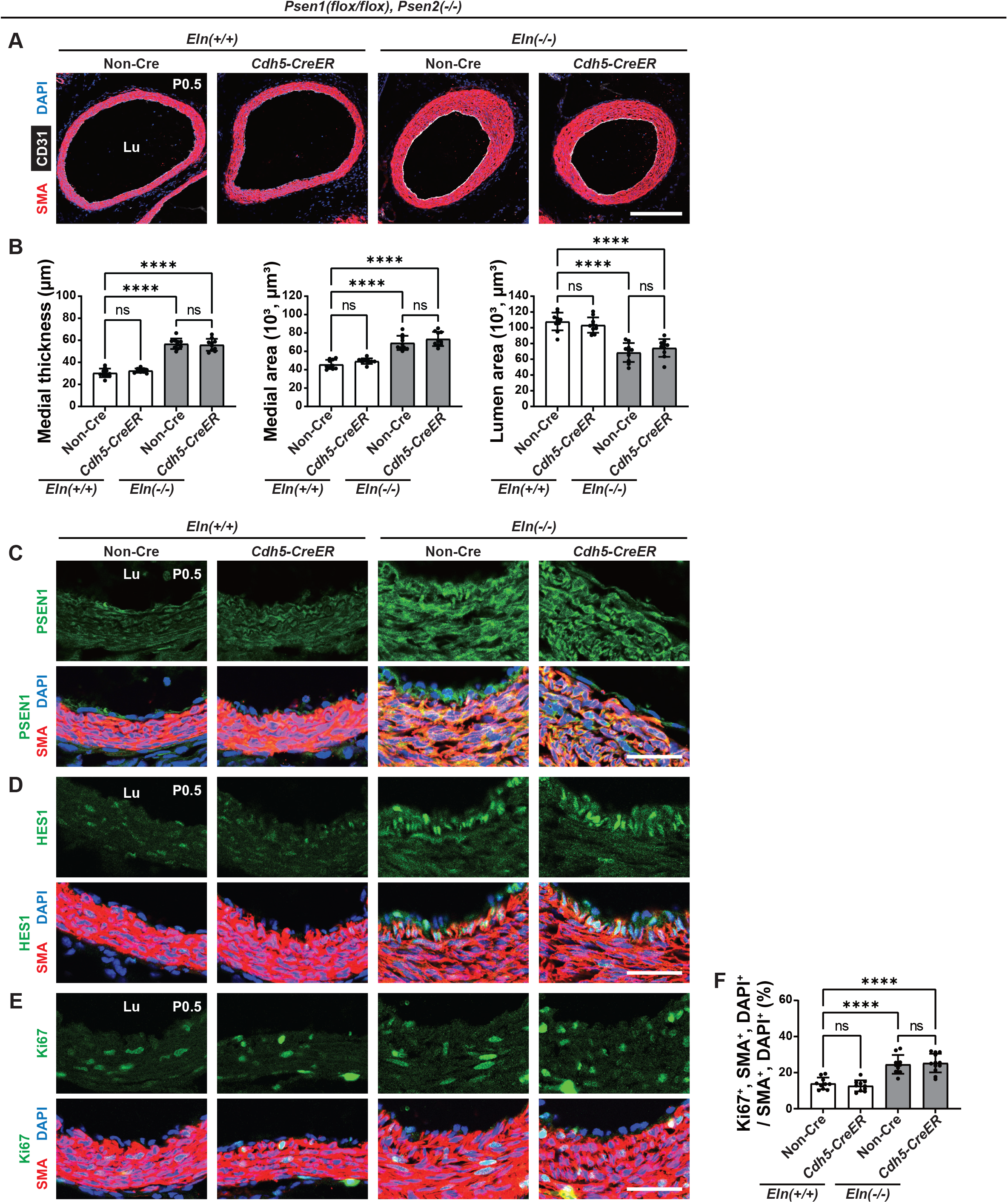
On the *Eln(-/-), Psen2(-/-)* background, EC-specific *Psen1* deletion does not attenuate hypermuscularization and stenosis. Transverse sections of the ascending aorta (cranial position, Fig. S1) from pups of indicated genotype at P0.5 were analyzed. **(A)** Sections were stained for SMA, CD31, and nuclei (DAPI). **(B)** Histograms represent aortic medial wall thickness, medial area and lumen area from sections as shown in **A**. n=10 mice. **(C-E)** Sections were stained for SMA, nuclei (DAPI), and either PSEN1 in **C**, HES1 in **D**, or Ki67 in **E**. **(F)** Histogram represents percent of SMCs that are Ki67^+^ in sections as in **E**. n=10-12 mice. Lu, lumen. ns, not significant. \*\*\*\**P* < 0.0001 by multifactor ANOVA with Tukey’s *post hoc* test. Scale bars, 200 μm (**A**) and 40 μm (**C-E**).

**Figure S7.**
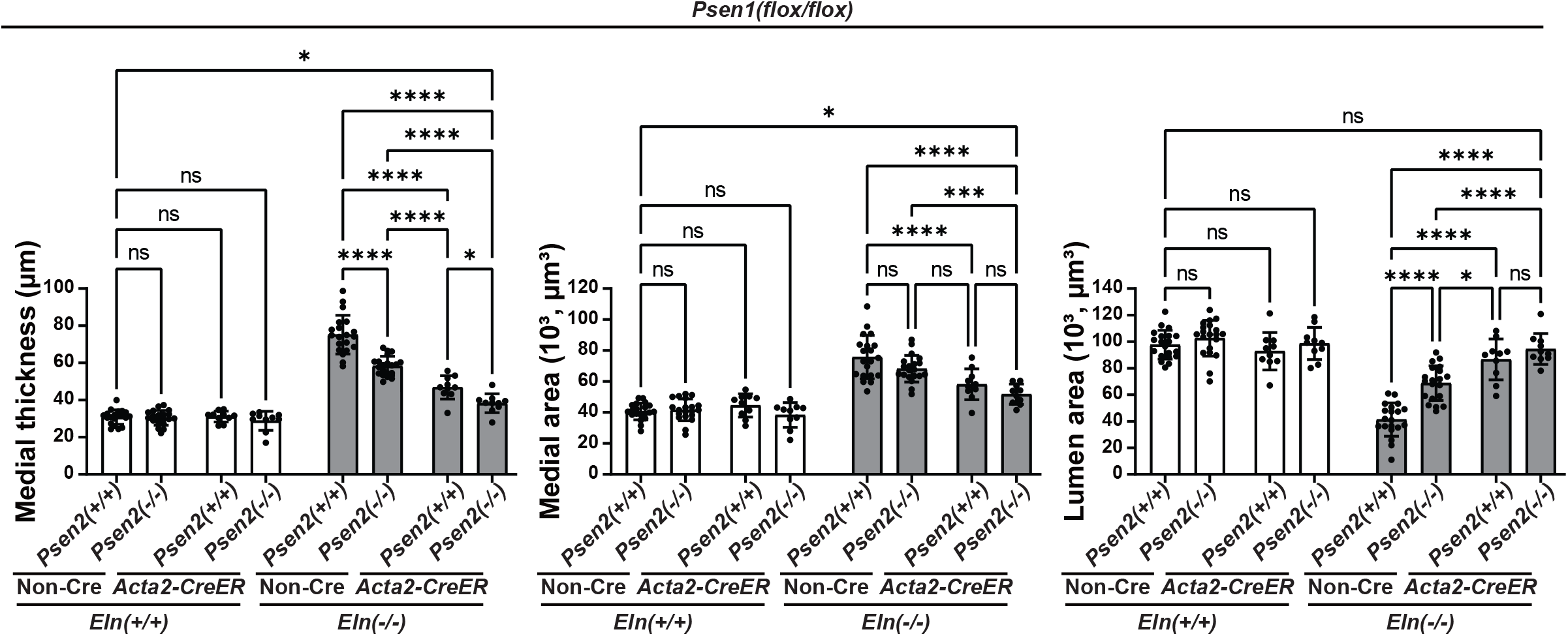
Combined deletion of *Psen1* in SMCs and *Psen2* globally has additional rescue effect on ascending aorta morphology in *Eln(-/-)* mice. (A) Histograms demonstrate combined data of ascending aorta medial wall thickness, medial area and lumen area from Figures. 2B, 3B, and 4B. n=10-20 mice.

**Figure S8.**
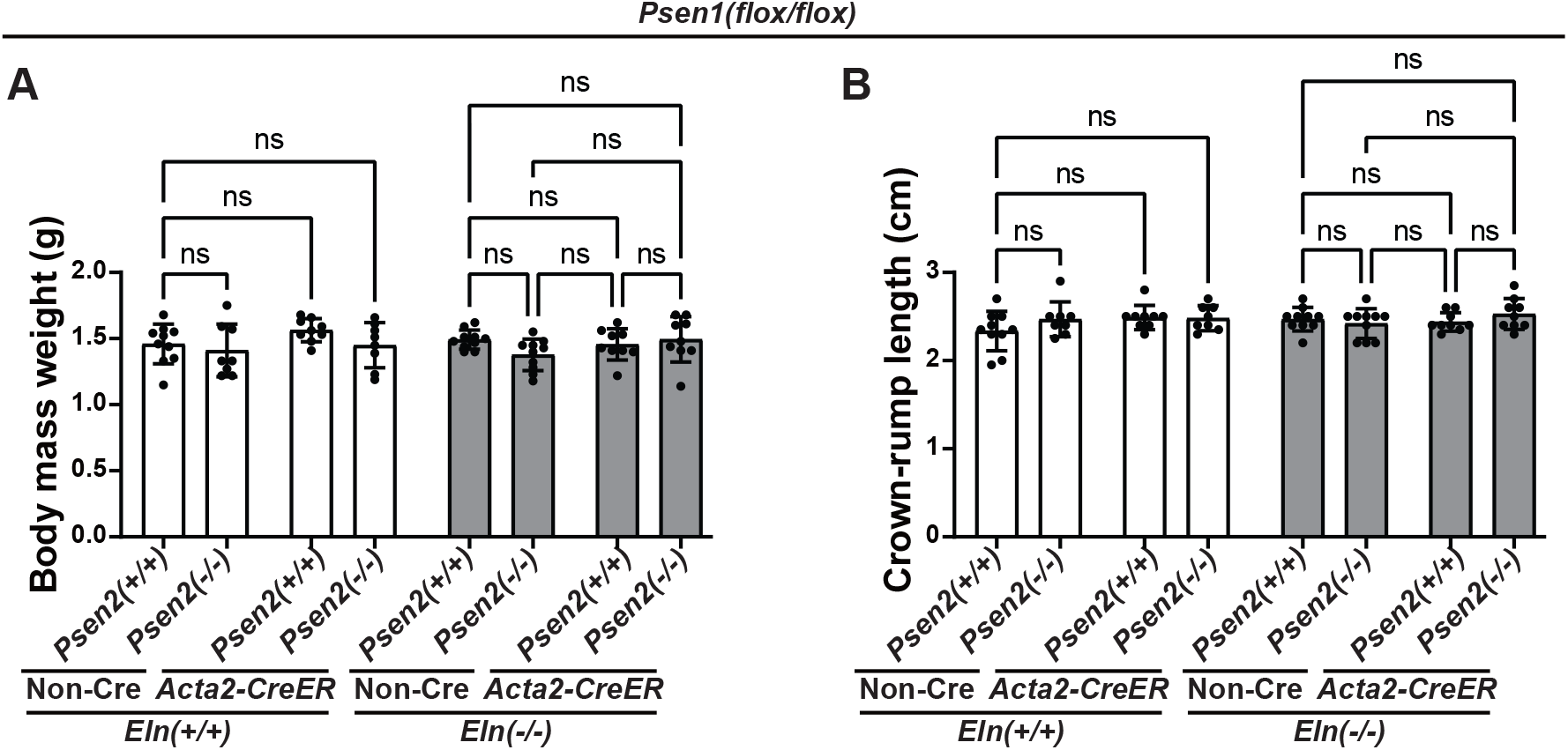
Deletion of *Psen1* in SMCs and/or *Psen2* globally does not alter body size. Histograms represent body mass weight (**A**) and crown-rump length (**B**) of each genotype. n=8-10 mice. ns, not significant. Multifactor ANOVA with Tukey’s *post hoc* test was used.

**Supplemental Table 1.**
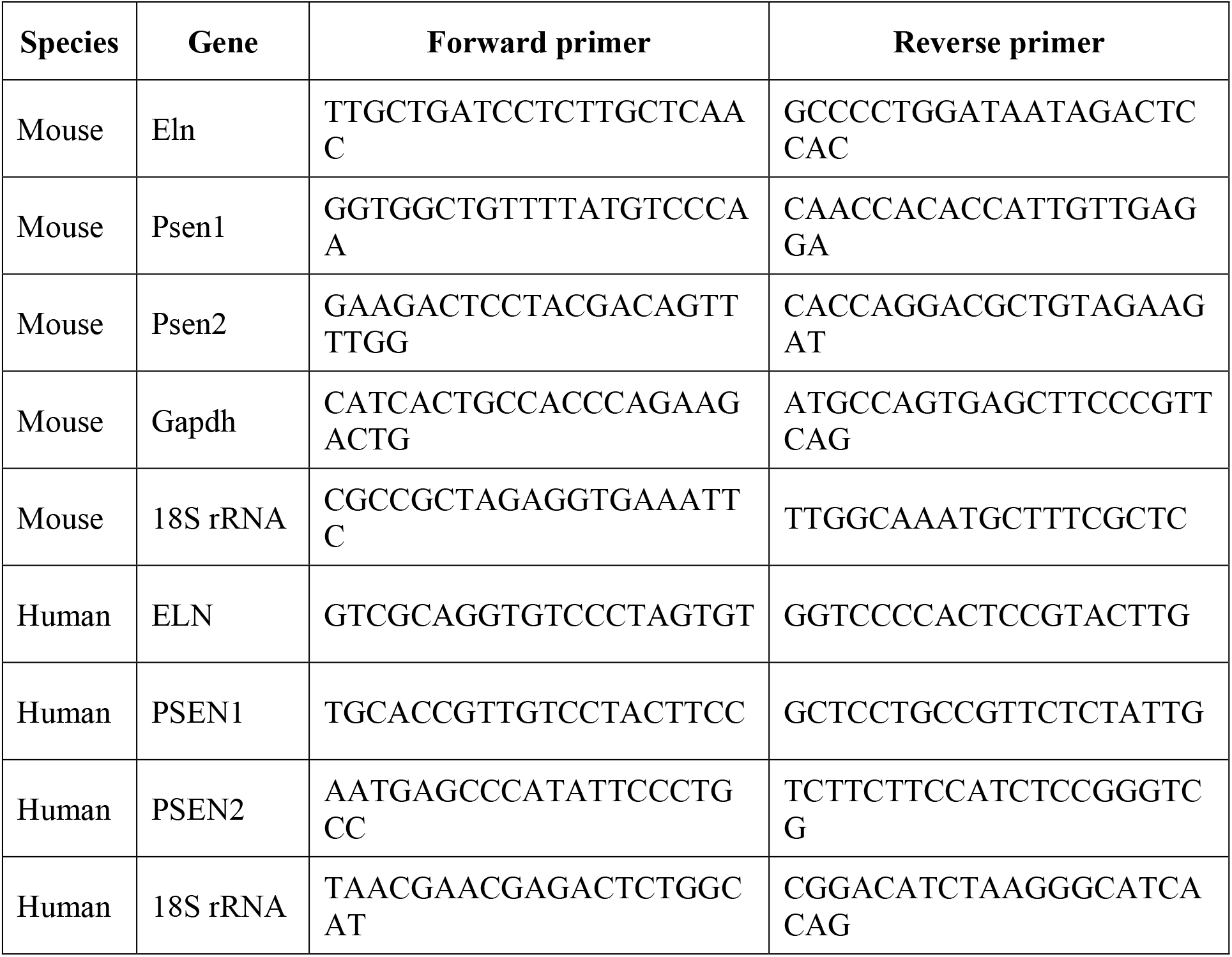
Primer pair sequences used for quantitative reverse transcription polymerase chain reactions.

**Supplemental Table 2.**
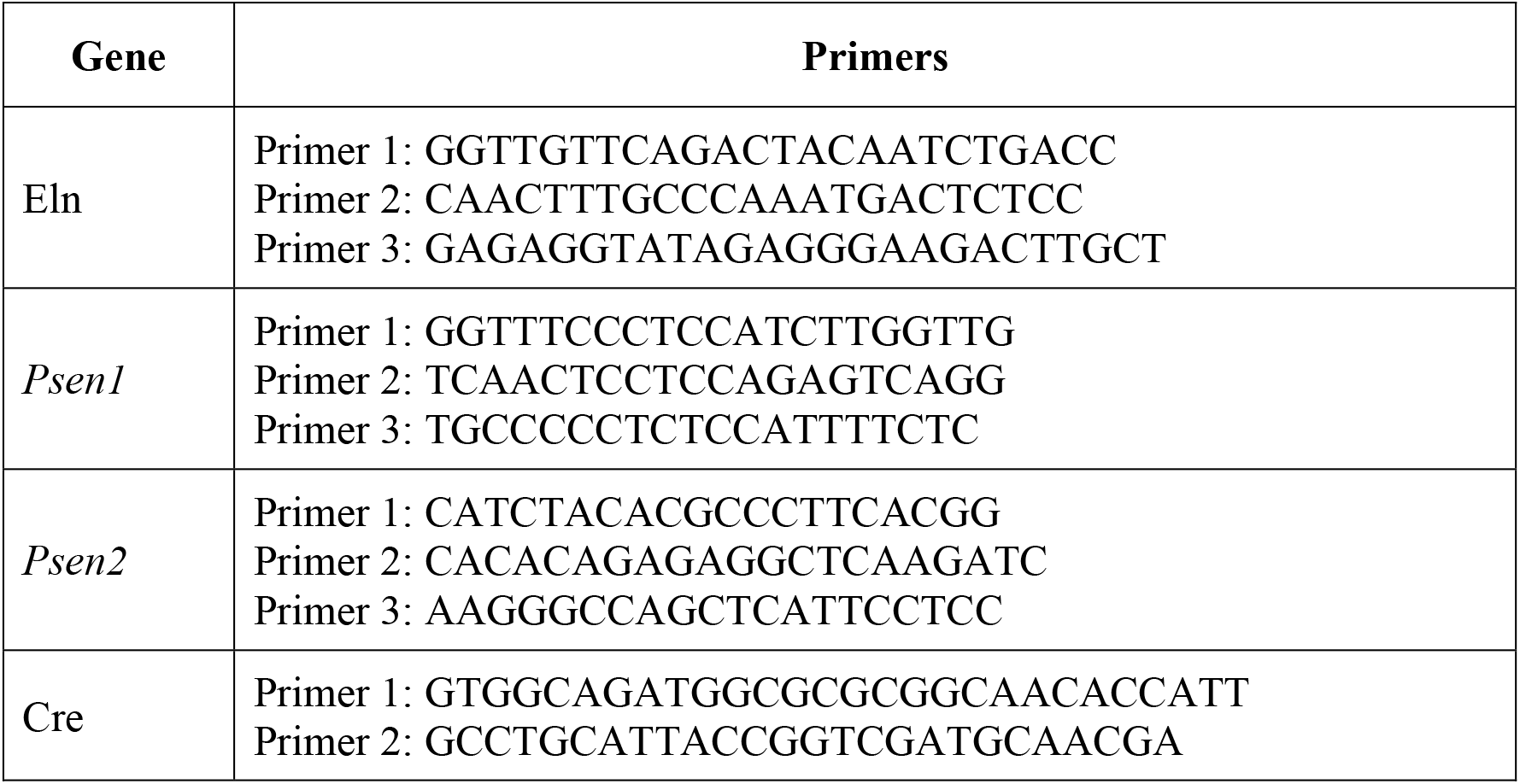
Primer sequences used for genotyping.

